# GeoDiver: Differential Gene Expression Analysis & Gene-Set Analysis For GEO Datasets

**DOI:** 10.1101/127753

**Authors:** Ismail Moghul, Suresh Hewapathirana, Nazrath Nawaz, Anisatu Rashid, Marian Priebe, Bruno Vieira, Fabrizio Smeraldi, Conrad Bessant

## Abstract

**Summary:** GeoDiver is an online web application for performing Differential Gene Expression Analysis (DGEA) and Generally Applicable Gene-set Enrichment Analysis (GAGE) on gene expression datasets from the publicly available Gene Expression Omnibus (GEO). The output produced includes numerous high quality interactive graphics, allowing users to easily explore and examine complex datasets instantly. Furthermore, the results produced can be reviewed at a later date and shared with collaborators.

**Availability:** GeoDiver is freely available online at http://www.geodiver.co.uk. The source code is available on Github: https://github.com/GeoDiver/GeoDiver and a docker image is available for easy installation.

## 1 Introduction

Gene expression analysis is a powerful methodology by which differences in gene expression profiles can be identified between sample populations. As such, these analyses are routinely used in a variety of research areas including cancer research (e.g. Arpino et al., 2013), other disease research (e.g. Emilsson et al., 2008), and drug development (e.g. Bai et al., 2013).

The recent exponential decrease in sequencing cost and advancement in microarray technology has resulted in an accumulation of large gene expression datasets; many of which are publicly available on the Gene Expression Omnibus (GEO) (Barrett et al., 2013). Despite having access to numerous large datasets, analysing them can be challenging and time-consuming.

Typical analyses on gene-expression data includes Differential Gene Expression Analysis (DGEA) and Gene-Set Analysis (GSA), which are used to find statistically significant differences in expression of specific genes or gene-sets between sample populations. There are a wide variety of R packages that can be used to analyse gene expression data (e.g. Robinson et al., 2010, Ritchie et al., 2015). However, the use of these R packages and the subsequent generation of visualisations requires coding proficiency, for example, in the R programming language.

Existing tools, such as the Geo2R (Barrett et al., 2013) and GSEA (Subramanian et al., 2005), can be used to carry out gene expression analysis, but do not produce extensive graphics for thorough visual analysis.

GeoDiver is an online web application that analyses existing GEO Datasets and GEO Data Series using DGEA and GSA. Interactive graphics are produced to visualise the results of these analyses as well as for the exploratory data analysis. These high-quality graphics include interactive (two- and three-dimensional) principal component analysis plots, heatmaps, interaction networks and volcano plots (see Figure 1). Users can also optionally login in order to store their results online, review past analyses and share these via a URL.

**Fig. 1.**
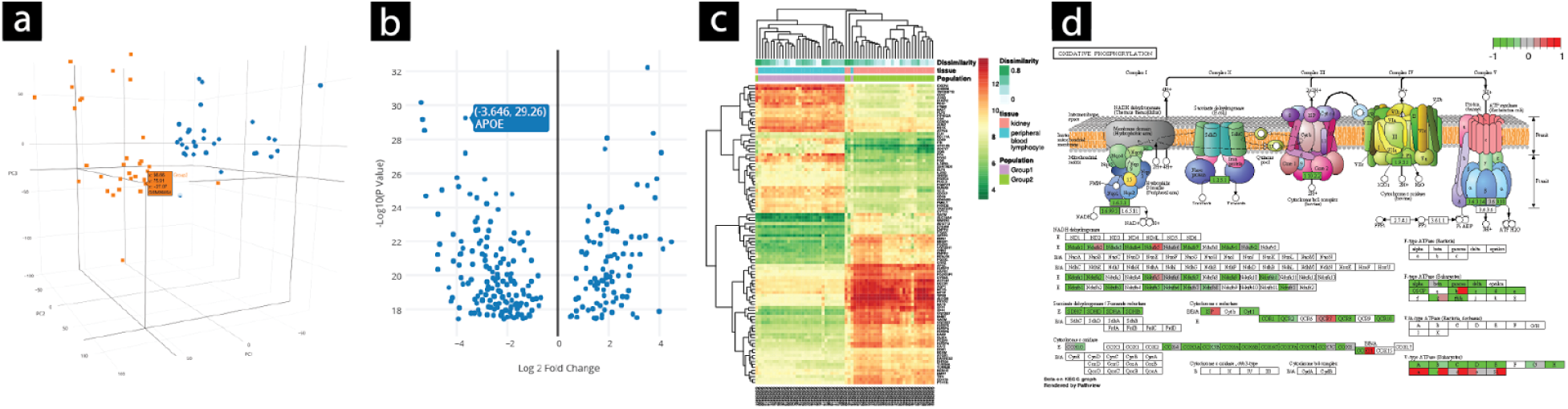
Exemplar GeoDiver Visualisations: a) An Interactive 3D PCA plot showing data points for the first three components, b) an interactive volcano plot showing significant genes c) a heatmap of the top differentially expressed genes and d) a colour-coded interaction network (oxidative phosphorylation). The above exemplar visualisations have been produced with GEO dataset GDS724, comparing Peripheral Blood Lymphocyte (as Group A) and Kidney (as Group B) expression levels.

## 2 Approach

In recent years a common workflow for analysing and reporting differential gene expression experiments has evolved, and it is this that has been implemented in GeoDiver.

In the first instance, users are required to enter the accession number of the GEO Dataset or Data Series to be analysed. GeoDiver downloads the dataset using the NCBI module from Bionode (bionode.io). Upon importing the data, GeoDiver initially carries out a preliminary analysis to determine whether the data needs to be scaled and automatically log transforms the data if necessary. GeoDiver additionally examines the data and imputes any missing values using KNN imputation. A principal component analysis is carried out to reveal any discrete clusters in the data. Moreover, the individual samples are analysed to determine the likelihood of a sample being an outlier.

### 2.1 Differential Gene Expression Analysis

GeoDiver uses the limma R package (Ritchie et al., 2015) to identify differentially expressed genes by fitting a linear model to each gene which estimates the fold change in expression while accounting for standard errors by applying empirical Bayes smoothing. Genes are ordered according to the fold change in the expression values.

This information is presented as an interactive table, a heatmap and a volcano plot. Upon clicking on a gene, users are also provided with an interactive bar chart displaying the gene expression levels for each sample expressing the gene. The Volcano plot has added interactivity showing the gene name, fold-change and p-value of each data point.

### 2.2 Generally Applicable Gene-Set Enrichment Analysis

Generally Applicable Gene-set Enrichment for Pathway Analysis (GAGE) is a variation of gene set enrichment analysis. Instead of sample randomization, it uses gene randomization, allowing it to carry out accurate analyses of datasets with fewer samples (Luo et al., 2009).

GeoDiver utilises the KEGG (Kanehisa and Goto, 2000) and Gene Ontology (Gene Ontology Consortium, 2004) databases to identify the pathways each gene in the data is associated with. Using this information, GeoDiver is able to identify the pathways that are significantly differentially expressed between the sample populations selected. This information is presented as an interactive table and a heatmap. Additionally, a colour-coded interaction network is also produced for analyses based on the KEGG database.

## 3 Usage & Implementation

GeoDiver provides users with a minimalistic graphical interface designed in accordance with Google’s Material Design Specifications.

Upon entering a GEO accession number, users can select the sample populations they wish to compare and immediately start their analysis. Users are also provided with several options to customise and fine-tune their analysis by being able to change over 20 parameters including the protocols used for clustering and false discovery rates.

A standard analysis takes a few minutes to complete and the results are displayed on the same page. Amongst the results produced for each analysis are high-quality vector-based visualisations, tab delimited tables and R data objects; all of which can be downloaded. GeoDiver’s web application has been written in the Ruby programming language (largely based on GeneValidator (Dragan et al., 2016)) while the DGEA and GAGE analyses are in R (see supplementary figure 1).

## 4 Discussion

Without adjusting any advanced parameters, users can easily run both DGEA and GSA within a few minutes of accessing the GeoDiver web application. For users who are familiar with gene-expression data, the option to adapt the advanced parameters of the analyses mirrors the flexibility that one might expect from writing a custom analysis script.

GeoDiver allows researchers to fully take advantage of the growing GEO resource and perform powerful analyses without the need for downloading or installing additional software, learning command line skills or having prior programming knowledge.

Several of the graphics produced are interactive; this helps users to interpret and understand the data they have analysed. Other graphics such as the heatmaps are high-resolution, information rich and can be easily exported for use in publications. GeoDiver is therefore not only designed to facilitate the analysis of gene-expression data but also to ensure that users are able to fully explore the results of their analysis. This is important as the ability to use such powerful analytical tools has to be paired with the corresponding ability to interpret the output.

